# Does Tesla valve work for microscale active swimmers? – a computational study

**DOI:** 10.1101/2022.03.31.486569

**Authors:** Ariel Rogers, Yong Wang

## Abstract

Although the Tesla valve is well-known for its diodicity for fluid flows and pressure drops, it is not clear whether and how the resistances experienced by active swimmers in the forward and reverse directions are different. Here we carried out a computational study on the diodicity of the Tesla valve for active swimmers in the absence of fluid flows. We simulated the active Brownian motion of the swimmers in a Tesla valve, followed by examining their trajectories and quantifying the fraction of active swimmers reaching the left or right end of the Tesla valve (in the forward or reverse direction, respectively). We also estimated the first passage time of the swimmers reaching the valve ends. We confirmed that, in the absence of fluid flows, the Tesla valve shows much higher resistance to active swimmers in the reverse direction than the forward direction. The current study provides a better understanding of the interaction of the Tesla valve with active swimmers and gives insight into potential applications of the Tesla valve in the filtering and sorting of motile microbes.

## 1. INTRODUCTION

The capability of controlling the flow of fluid, gas, particles, and microbes is important in various applications[1, 2]. Such capability is mostly achieved based on reciprocating devices with moving parts, while fixed or no-moving-parts devices and valves have been proposed and studied[3]. Fixed devices/valves are attractive due to their simplicity of fabrication and maintenance[3]. One example of fixed valves is the Tesla valve, invented by Nikola Tesla[4] in 1920. The Tesla valve consists of linked, looped, and repeated lanes, and is diodic for fluid flows and pressure drops; in other words, it allows fluid to pass through in one direction (the forward direction) but exerts significant resistance in the other direction (the reverse direction)[4–6]. In the past years, the Tesla valve has attracted extensive computational and experimental studies, ranging from understanding its operations, to optimizing its performance, developing novel micro-pumps, and designing hybrid jet actuators[5–12].

Previous studies have focused on the Tesla valve for flows of fluids or gases that are associated with pressure drops[5–14]. However, the effectiveness of the Tesla valve in the absence of flows or pressure drops was rarely studied. More importantly, investigations on how the active matters interact with the Tesla valve have been seldom reported; and thus many questions remain unexplored. For example, it is unclear how diodic the Tesla valve is for active swimmers without fluid flows (or pressure drops). Do active swimmers feel different resistances in the forward direction of the Tesla valve compared to the reverse direction? A better understanding of the diodicity of the Tesla valve for active swimmers, as well as the interaction of the Tesla valve with active matters in general, is expected to provide insight into understanding the function of certain biological structures, such as shark spiral intestines[15], and may pave a way to developing new applications of the Tesla valve in microbiological research and clinical settings[16–18].

In this work, we carried out a computational study on the diodicity of the Tesla valve for active swimmers in the absence of fluid flows. We simulated the active Brownian motion[19] of the swimmers in a Tesla valve, followed by examining their trajectories and quantifying the percentage of active swimmers reaching the left or right end (in the forward or reverse direction) of the Tesla valve. We also estimated the first passage time of the swimmers reaching the ends of the Tesla valve. This computational study confirmed the diodicity of the Tesla valve for active swimmers even in the absence of fluid flows.

## II. MATERIALS AND METHODS

### A. Geometry of the Tesla valve used in this study

The geometry of the Tesla valve used in this study is shown in Fig. 1A. The design is similar to the original one by Nikola Tesla[4], while the dimensions were reduced to micro-scale. The width of the looped lanes of the Tesla valve was 4 *µ*m, and the number of repeating loops was 9. The Tesla valve was connected to two circular pools at both ends, with diameters of 40 *µ*m.

**FIG. 1.**
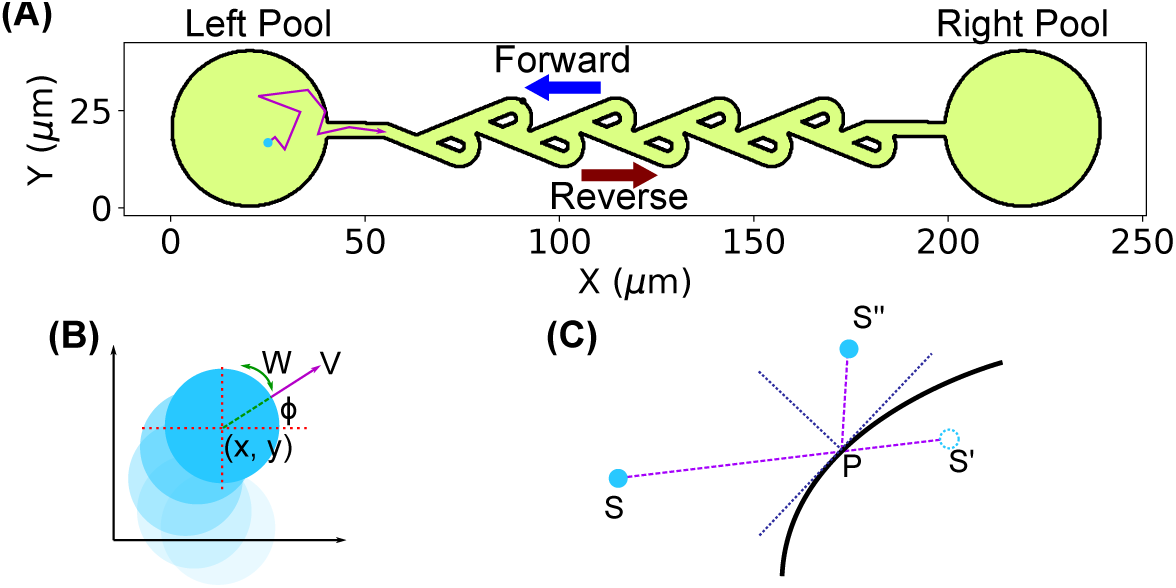
Geometry of a microscale Tesla valve and simulation of active swimmers. (**A**) The geometry of the microscale Tesla valve in this study. The forward and reverse directions of the Tesla valve were indicated by the blue and red arrows, respectively. (**B**) Active swimmer described by its location (*x, y*) and orientation *φ*, moving with a deterministic linear velocity *V*, a deterministic angular velocity *W*, and random translational and rotational diffusion. (**C**) Reflection of an active swimmer at the surface of the Tesla valve. *S*: original position; *S*^*/*^: tentative position after a move; *S*^*//*^: position after reflection.

### B. Simulation of the motion of active swimmers in the Tesla valve

Simulation of the motion of active swimmers were performed following Volpe et al.[19], which was previously used by us for understanding the interaction of bacteria with micro-pillars[20]. Briefly, an active swimmer is placed at an initial location and modeled as a sphere of radius *R* = 1 *µ*m, described by its location (*x, y*) and orientation *θ*. The motion of the active swimmer is due to a self-propelling force (leading to a deterministic, directional linear velocity *V*), a torque (leading to a deterministic, rotational angular velocity *W*), and random diffusion (both translational and rotational), as shown in Fig. 1B[19]. In the simulations of an active swimmer, its tentative orientation *θ*_*i*_ and location (*x*_*i*_, *y*_*i*_) at time step *i* were first calculated from the previous step: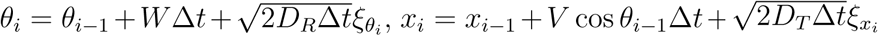, and 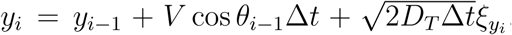, where Δ*t* is the time step size, *D*_*R*_ = *k*_*B*_*T/*8*πηR*^3^ is the rotational diffusion coefficient, *D*_*T*_ = *k*_*B*_*T/*6*πηR* is the translational diffusion coefficient, *η* = 10^*−*3^ Pa*·*s is the viscosity of the fluid, *k*_*B*_ is the Boltzmann constant, *T* is the temperature, *ξ*’s are random numbers from a Gaussian distribution with a mean of zero and a standard deviation of one[19, 20]. If the active swimmer did not collide with the surfaces of the Tesla valve, the tentative values at step *i* were taken; however, if the active swimmer collided with the surface, the tentative bacterial location (*x*_*i*_, *y*_*i*_) was reflected at the boundary of the surface to the location at step *i* as described in Volpe et al.[19] (Fig. 1C). For each simulation, 100000 steps with a step size of Δ*t* = 0.02 s were performed, and the locations of the active swimmers were recorded. Simulations with different deterministic linear velocities (*V* = 5, 10, 15, 20 *µ*m/s), deterministic angular velocities (*W* = 0, 0.1, 0.5, 1.0 rad/s), and initial positions: (*x*_0_, *y*_0_) = (20, 20) *µ*m (the center of the left pool), (*x*_0_, *y*_0_) = (220, 20) *µ*m (the center of the right pool), and (*x*_0_, *y*_0_) = (130, 20) *µ*m (the center of the Tesla valve). To estimate the statistics, 96 repeated simulations were run for each set of parameters.

### C. Simulation of the motion of passive particles in the Tesla valve

The simulations for passive particles were performed the same as the active swimmers by setting *V* = 0 *µ*m/s and *W* = 0 rad/s; in addition, a larger step size (Δ*t* = 0.1 ms) and a longer simulation time (1000000 steps) were used.

## III. RESULTS AND DISCUSSIONS

With the Tesla valve sketched in Fig. 1A, we simulated the motion of active swimmers starting at three different positions: the center of the left pool 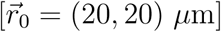, the center of the right pool 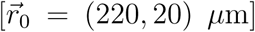, and the center of the Tesla valve 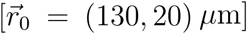. Two examples of trajectories for each starting location are shown in Fig. 2, with deterministic linear and angular velocities of *V* = 15 *µ*m/s and *W* = 0 rad/s. Active swimmers that started at the left pool did not reach the right pool (in the reverse direction of the Tesla valve), although attempts of entering the looped lanes of the Tesla valve were made (Fig. 2A). In contrast, active swimmers that started at the right pool reached the left pool (in the forward direction of the Tesla valve) and were then “trapped” in the left (Fig. 2B). In addition, active swimmers that started at the center of the Tesla valve reached the left pool while a few attempts to the right were observed (Fig. 2C). These observations suggest that, even if fluid flows were absent, the Tesla valve appeared diodic to the active swimmers: they experienced higher resistance in the reverse direction (left-to-right) of the Tesla valve than in the forward direction (right-to-left).

**FIG. 2.**
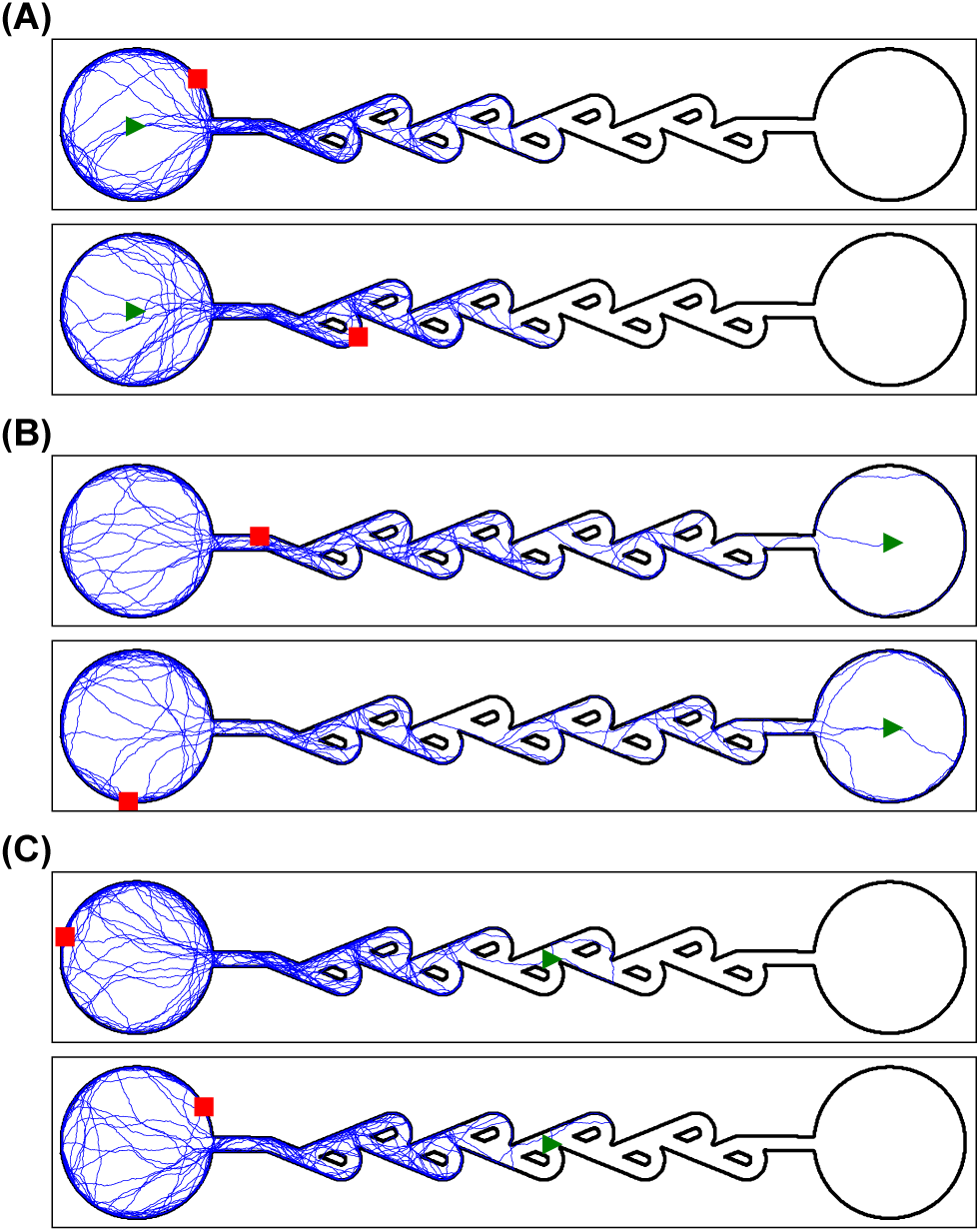
Examples of trajectories of individual active swimmers (*V* = 15 *µ*m/s, *W* = 0 rad/s) with initial positions (**A**) at the center of the left pool, (**B**) at the center of the right pool, and (**C**) at the center of the Tesla valve. The initial and final positions of the active swimmers were indicated by the green triangles and red squares, respectively.

To systematically examine the diodicity of the Tesla valve for active swimmers in the absence of fluid flows, we varied the deterministic linear and angular velocities (*V* = 5, 10, 15, and 20 *µ*m/s; *W* = 0, 0.1, 0.5, and 1.0 rad/s) and quantified the percentage (*P*_*a*_) of active swimmers that reached the left (*R → L* and *C → L*) and the right (*L → R* and *C → R*) pools. We observed that nearly 100% of the active swimmers reached the left pool for all the different velocities (*R → L* and *C → L*, Fig. 3) at *W* = 0 rad/s. In contrast, with the same parameters (i.e., *V* and *W*), none or only a small fraction reached the right pool (*L → R* and *C → R*, Fig. 3). It was also observed that, at lower deterministic linear velocities, the fraction of active swimmers reaching the right pool (in the reverse direction of the Tesla valve) were higher (e.g., *V* = 5 *vs*. 20 *µ*m/s, Fig. 3), which implies that the diodicity of the Tesla valve may diminish for passive particles (*V →* 0) in the absence of fluid flows, which was further investigated below.

**FIG. 3.**
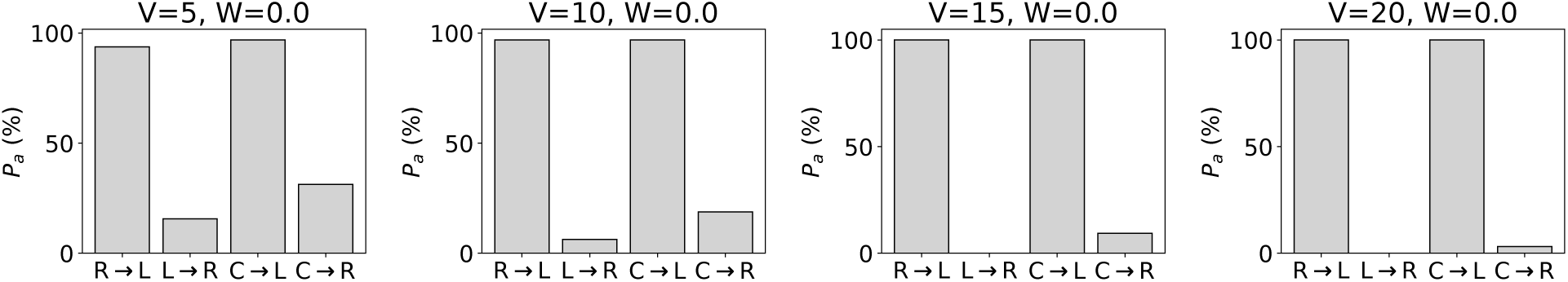
The percentage of active swimmers (*P*_*a*_) reaching the left or right pool with different initial positions and deterministic linear velocities (*V*, in *µ*m/s) but with zero angular velocity (*W* = 0 rad/s). *R → L*: starting at the right pool and reaching the left pool; *L → R*: starting at the left pool and reaching the right pool; *C → L*: starting at the center of the Tesla valve and reaching the left pool; *C → R*: starting at the center of the Tesla valve and reaching the right pool.

We observed that active swimmers with higher deterministic angular velocities experienced higher resistances in the forward direction of the Tesla valve. For example, at *V* = 15 *µ*m/s, the percentage of active swimmers reaching the left pool while started at the right pool reduced from 100% at *W* = 0 rad/s and *W* = 0.1 rad/s to 72% at *W* = 0.5 rad/s and 13% at *W* = 1.0 rad/s (SI Fig. 1). Similar trends were observed for other linear velocities (SI Fig. 1). This result is reasonable considering that the Tesla valve in the forward direction (right-to-left) is a straight, smooth path, but those swimmers with large angular velocities would deviate from the straight path and collide with the surfaces.

We then examined the percentage of active swimmers reaching the left (*P*_*L*_) or right (*P*_*R*_) pool as a function of time. As shown in Fig. 4 (for *W* = 0 rad/s), the percentage of active swimmers reaching the right pool (in the reverse direction of the Tesla valve) increased slowly at lower *V* and remained flat at higher *V* values (Fig. 4A and 4C). In contrast, the fraction of active swimmers reaching the left pool (in the forward direction of the Tesla valve) quickly increased to ∼100% for all the *V* values from 5 to 20 *µ*m/s (Fig. 4B and 4D). Similar results were observed for nonzero angular velocities: the resistance in the reverse direction remained much more significant for all angular velocities (SI Fig. 2). These results confirmed the diodicity of the Tesla valve for active swimmers in the absence of fluid flows.

**FIG. 4.**
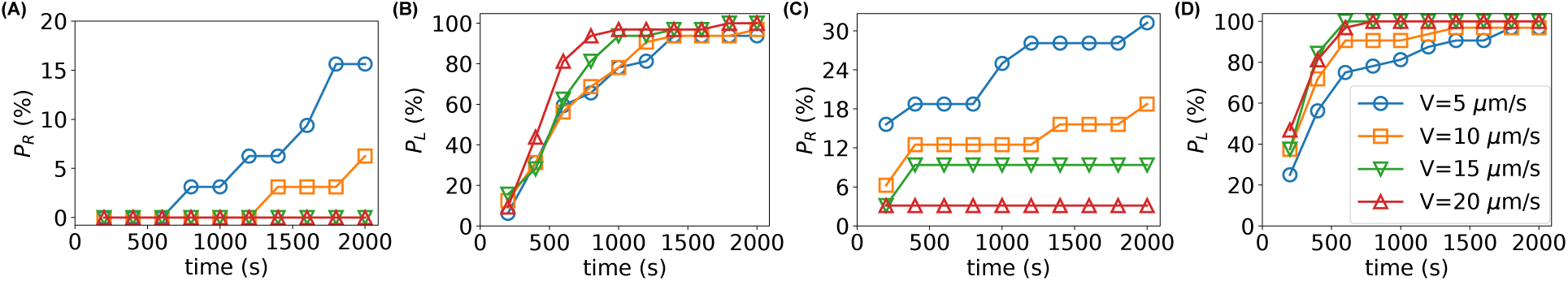
(**A, B**) The percentage of active swimmers starting at the (**A**) left or (**B**) right pool and reaching the opposite pool as a function of time, with different deterministic linear velocities (*V*, in *µ*m/s) and zero angular velocity (*W* = 0 rad/s). (**C, D**) The percentage of active swimmers starting at the center of the Tesla valve and reaching the (**C**) right or (**D**) left pool as a function of time, with different deterministic linear velocities (*V*, in *µ*m/s) and zero angular velocity (*W* = 0 rad/s).

As the first passage time is an important metric for understanding the dynamics of stochastic systems[21, 22], we investigated the first passage time *τ*_1_ of the active swimmers reaching the left or right pool (in the forward and reverse directions of the Tesla valve, respectively) when started at different initial positions. As shown in Fig. 5A (for *W* = 0 rad/s), the mean first passage times ⟨*τ*_1_⟩ of the active swimmers in the forward direction of Tesla valve (*R → L* and *C → L*) were 2–6 times shorter than those in the reverse direction (*L → R* and *C → R*). Similar results were observed for nonzero angular velocities (SI Fig. 3). We note that the mean first passage times in the reverse direction were underestimated because many of the active swimmers did not reach the right pool at the end of the simulation (2000 s, or 100000 steps), which was seen clearly in the distribution of the first passage time (Fig. 5B). For example, most or all of the active swimmers did not reach the right pool (in the reverse direction of the Tesla valve) at the end of the simulation of 2000 s when started at the left pool (LR/*P*_*R*_ in Fig. 5B). In contrast, the majority of the active swimmers reached the left pool from the right pool (in the forward direction of the Tesla valve) in less than 1000 s with peaks around 500 s (RL/*P*_*L*_ in Fig. 5B).

**FIG. 5.**
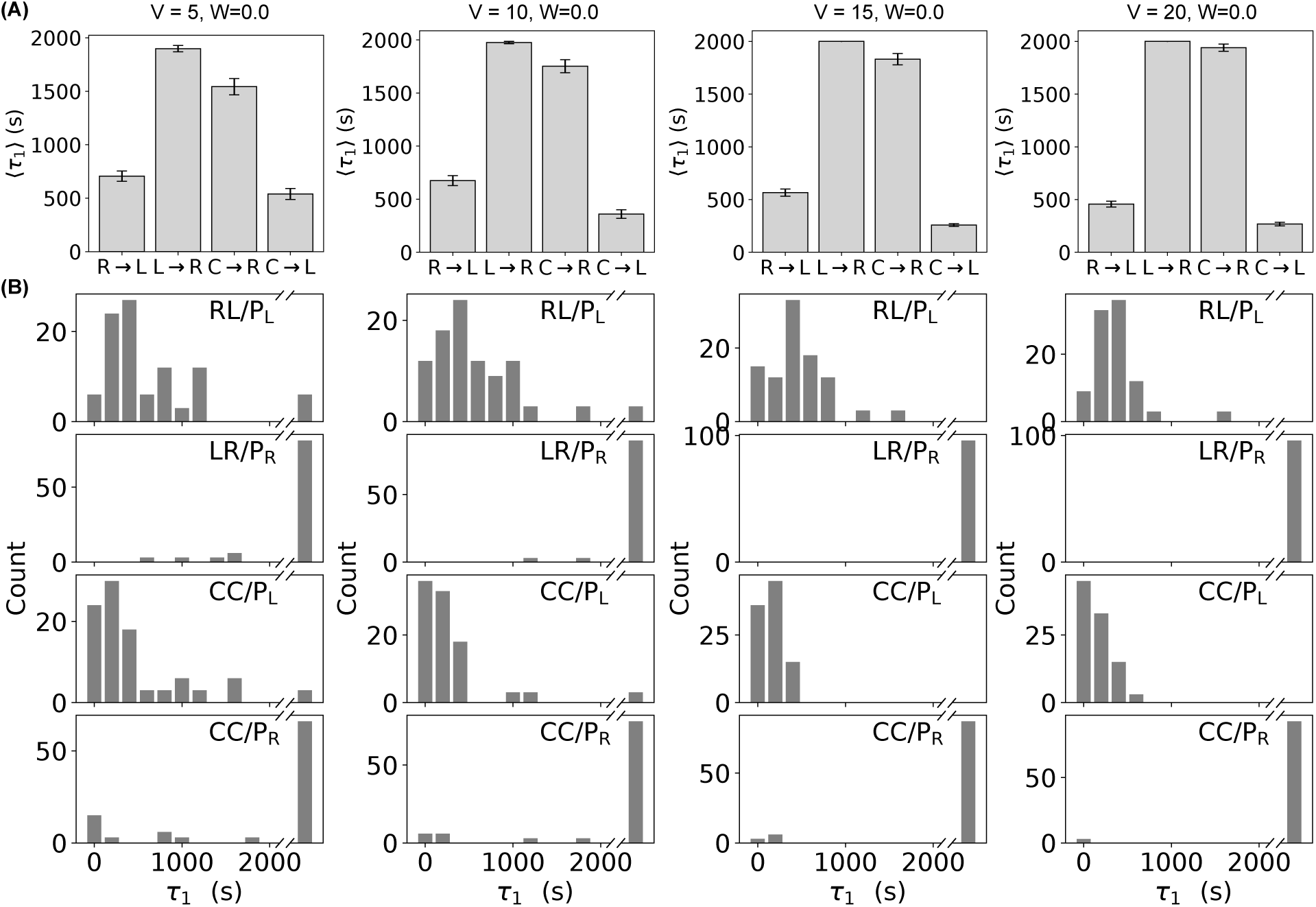
First passage time *τ*_1_ of active swimmers. (**A**) The average first passage time *(τ*_1_) of active swimmers reaching the left or right pool with different initial positions and deterministic linear velocities (*V*, in *µ*m/s), with zero angular velocity (*W* = 0 rad/s). (**B**) Distributions of the first passage time *τ*_1_ of active swimmers reaching the left or right pool with different initial positions and deterministic linear velocities (*V*, in *µ*m/s), with zero angular velocity (*W* = 0 rad/s). Line breaks indicate *>* 2000 s.

Lastly, we performed simulations for passive particles (*V* = 0 *µ*m/s and *W* = 0 rad/s), and confirmed that the diodicity of Tesla valve is much less significant for passive particles. For example, the percentage (*P*_*a*_) of passive particles reaching the left or right pool were similar (Fig. 6A): the difference in *P*_*a*_ was 100% for active swimmers with *V* = 15 *µ*m/s and *W* = 0 rad/s (*R → L vs. L → R*, Fig. 3); in contrast, the difference was only 5.7% for passive particles (*R → L vs. L → R*, Fig. 6A). In addition, the increase in *P*_*a*_ was similar in both directions for passive particles (RL/*P*_*L*_ *vs*. LR/*P*_*R*_, and CC/*P*_*L*_ *vs*. CC/*P*_*R*_, Fig. 6B).

**FIG. 6.**
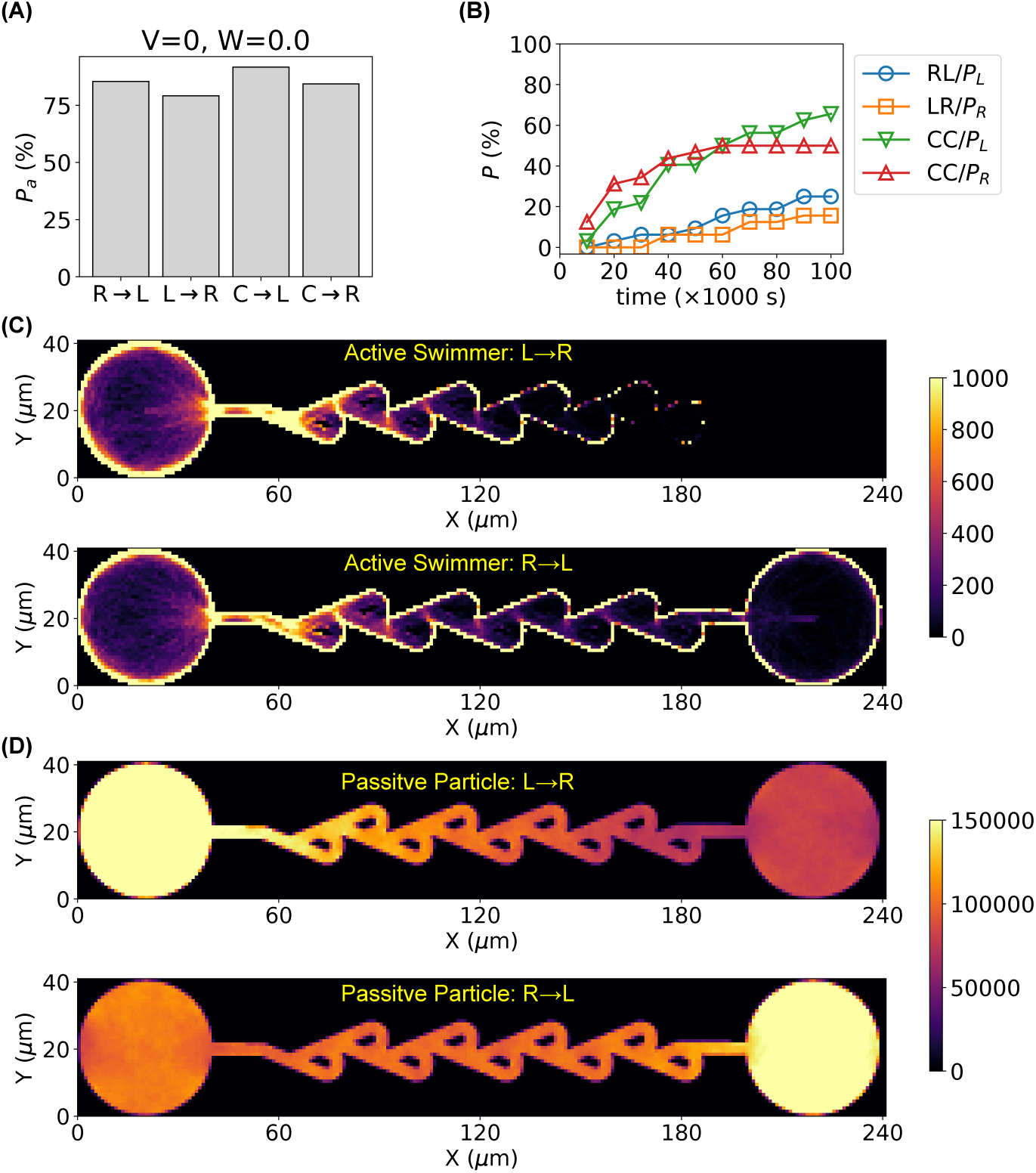
Simulated results for passive particles and comparison with active swimmers. (**A**) Percentage of passive particles arriving at the left or right pools with different initial positions. (**B**) Percentage of passive particles starting at different locations and arriving at different locations as a function of time. RL/*P*_*L*_: starting at the right pool and arriving at the left pool; LR/*P*_*R*_: starting at the left pool and arriving at the right pool; CC/*P*_*L*_: starting at the center of the Tesla valve and arriving at the left pool; CC/*P*_*R*_: starting at the center of the Tesla valve and arriving at the right pool. (**C, D**) Density maps for the positions of (**C**) active swimmers or (**D**) passive particles starting at the left pool (*L → R*) or the right pool (*R → L*).

To have a comprehensive view of the locations of the active swimmers and passive particles during the whole time span of the simulations, we visualized the 2D density of the swimmers and particles in Fig. 6C and 6D. The diodicity of the Tesla valve for active swimmers in the absence of fluid flows was further highlighted (Fig. 6C). Another observation is that, while the active swimmers tended to accumulate at the surfaces of the pools and the Tesla valve, similar to other micro-structures[19], the passive particles were more uniformly distributed.

## IV. CONCLUSIONS

To conclude, we computationally investigated whether and how the Tesla valve is diodic for active swimmers in the absence of fluid flows. The motion of active swimmers in the Tesla valve was simulated as active Brownian motion[19]. We examined the trajectories of the active swimmers that started at different locations of the Tesla valves (center and ends), and quantified the percentage of active swimmers reaching the left or right end (in the forward or reverse direction of the Tesla valve), as well as their first passage time to the ends. We found that active swimmers prefer the forward direction of the Tesla valve to the reverse direction, while such preference is significantly diminished for passive particles. This computational study confirmed the diodicity of the Tesla valve for active swimmers in the absence of fluid flows.

It would be interesting to experimentally verify the results of the current study. Motile bacteria (e.g., *E. coli*) would be potential candidates of active swimmers[20, 23, 24], while microscale Tesla valve could be fabricated using photolithography processes and/or microscale 3D printing[16, 20, 25, 26]. Such experimental investigations are expected to not only verify the results from the current study but also provide insight into the modeling of the active Brownian motion.

Studies have been reported for improving the performance and enhancing the diodicity of the Tesla valve for fluid flows based on optimization of the shape[7, 8, 12]. Although it is out of the scope of the current work, we expect that shape-based optimizations will improve the diodicity of the Tesla valve for active swimmers in the absence of fluid flows. In addition, the observed dependence of the Tesla valve’s diodicity on the properties of the active swimmers in this study suggests that the Tesla valve may be optimized against certain active swimmers when desired.

The current study also suggests that microscale Tesla valves may be of great use in filtering, chromatography, and sorting applications involving active swimmers such as motile microbes. For example, the microscale Tesla valve may be used to prevent motile microbes from escaping (along the reverse direction of the Tesla valve) while allowing nutrients/wastes to exchange freely. In addition, the current study suggested that the rate of active swimmers passing through the Tesla valve in the forward direction depends on both their linear velocity and angular velocity, which can be exploited in chromatography and/or sorting of a mixture of active swimmers with different properties.

## V. ACKNOWLEDGMENT

This work was supported by the National Science Foundation (Grant No. 1826642 and 2129225) and the Arkansas Biosciences Institute (Grants No. ABI-0189, ABI-0226, ABI-0277, ABI-0326, ABI-2021, and ABI-2022). We are also grateful for support from the Arkansas High-Performance Computing Center (AHPCC), which is funded in part by the National Science Foundation (grants no. 0722625, 0959124, 0963249, and 0918970) and the Arkansas Science and Technology Authority.

