## Supplementary material for "Does Tesla valve work for microscale active swimmers? – a computational study": SI

### Supplementary Information

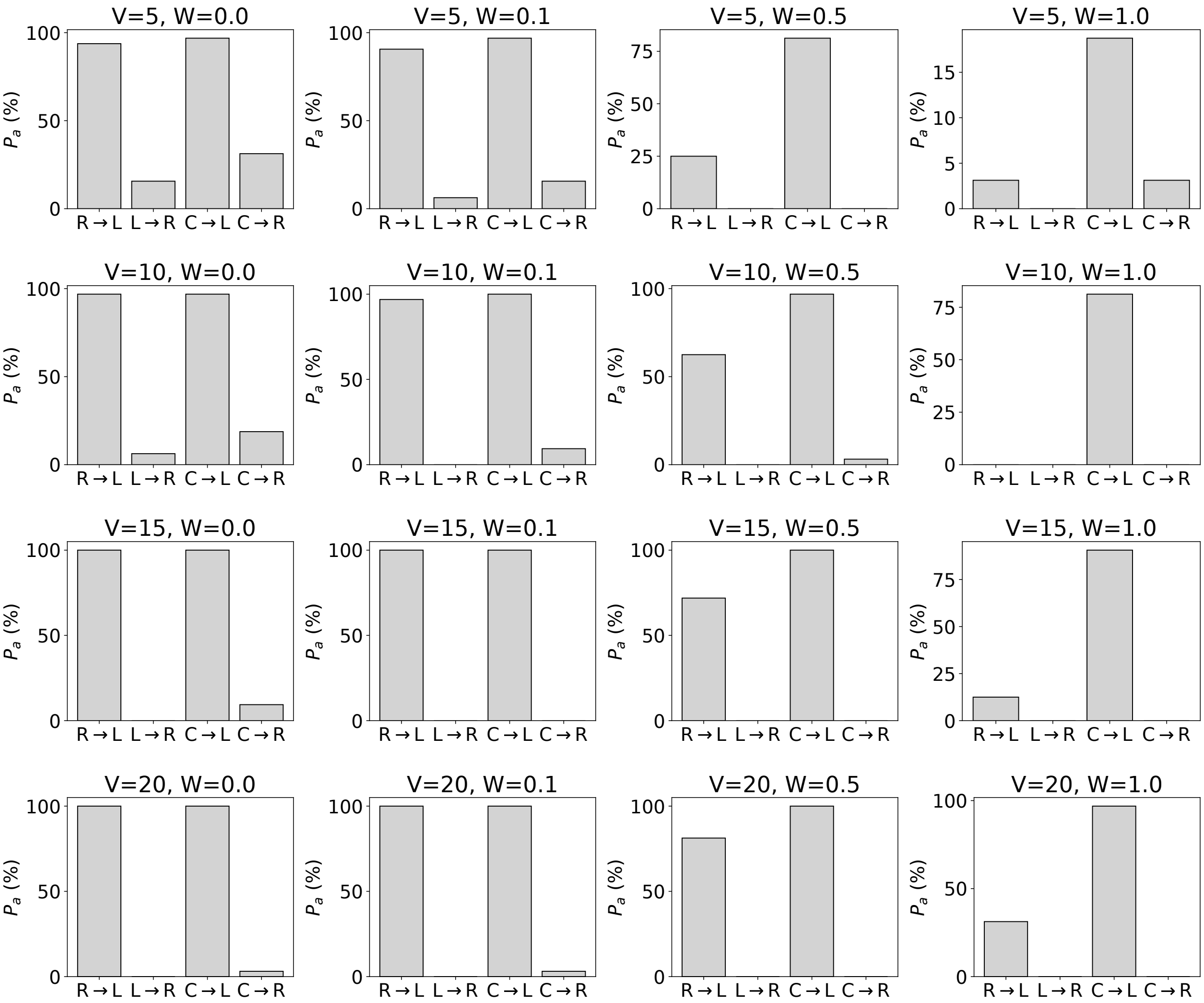


SI Figure 1: Percentage of active swimmers reaching the left or right pool with different initial positions, deterministic linear velocities ($V$, in $\mu$m/s), and deterministic angular velocities ($W$, in rad/s). $R \rightarrow L$: starting at the right pool and reaching the left pool (in the forward direction of the Tesla valve); $L \rightarrow R$: starting at the left pool and reaching the right pool (in the reverse direction of the Tesla valve); $C \rightarrow L$: starting at the center of the Tesla valve and reaching the left pool (in the forward direction of the Tesla valve); $C \rightarrow R$: starting at the center of the Tesla valve and reaching the right pool (in the reverse direction of the Tesla valve).

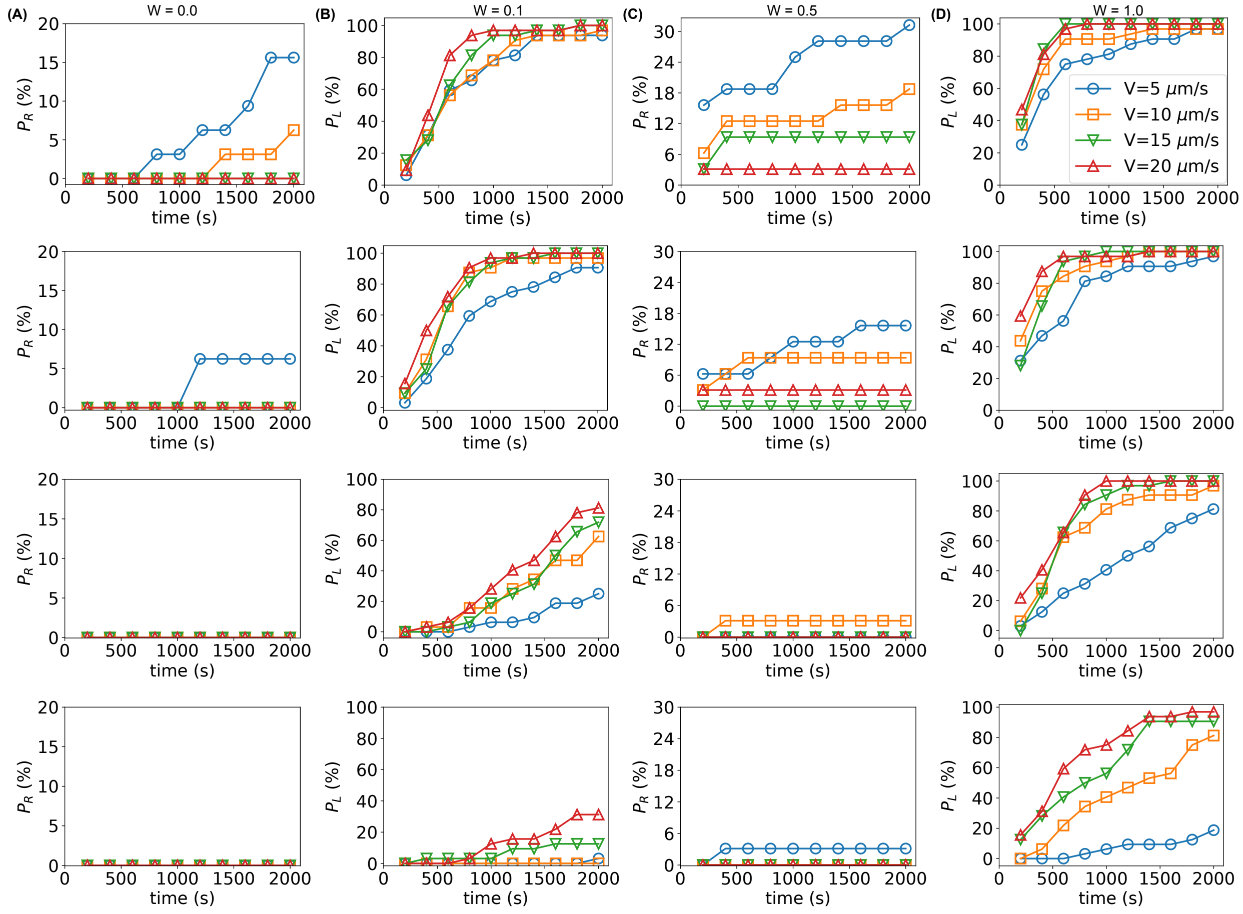


SI Figure 2: (**A**, **B**) Percentage of active swimmers starting at the (column **A**) left or (column **B**) right pool and reaching the opposite pools as function of time, with different deterministic linear velocies ($V$, in $\mu$m/s) and angular velocities ($W$, in rad/s). (**C**, **D**) Percentage of active swimmers starting at the center of the Tesla valve and arriving at the (column **C**) right or (column **D**) left pools as function of time, with different deterministic linear velocies ($V$, in $\mu$m/s) and angular velocities ($W$, in rad/s). Different symbols/colors indicate different deterministic linear velocities.

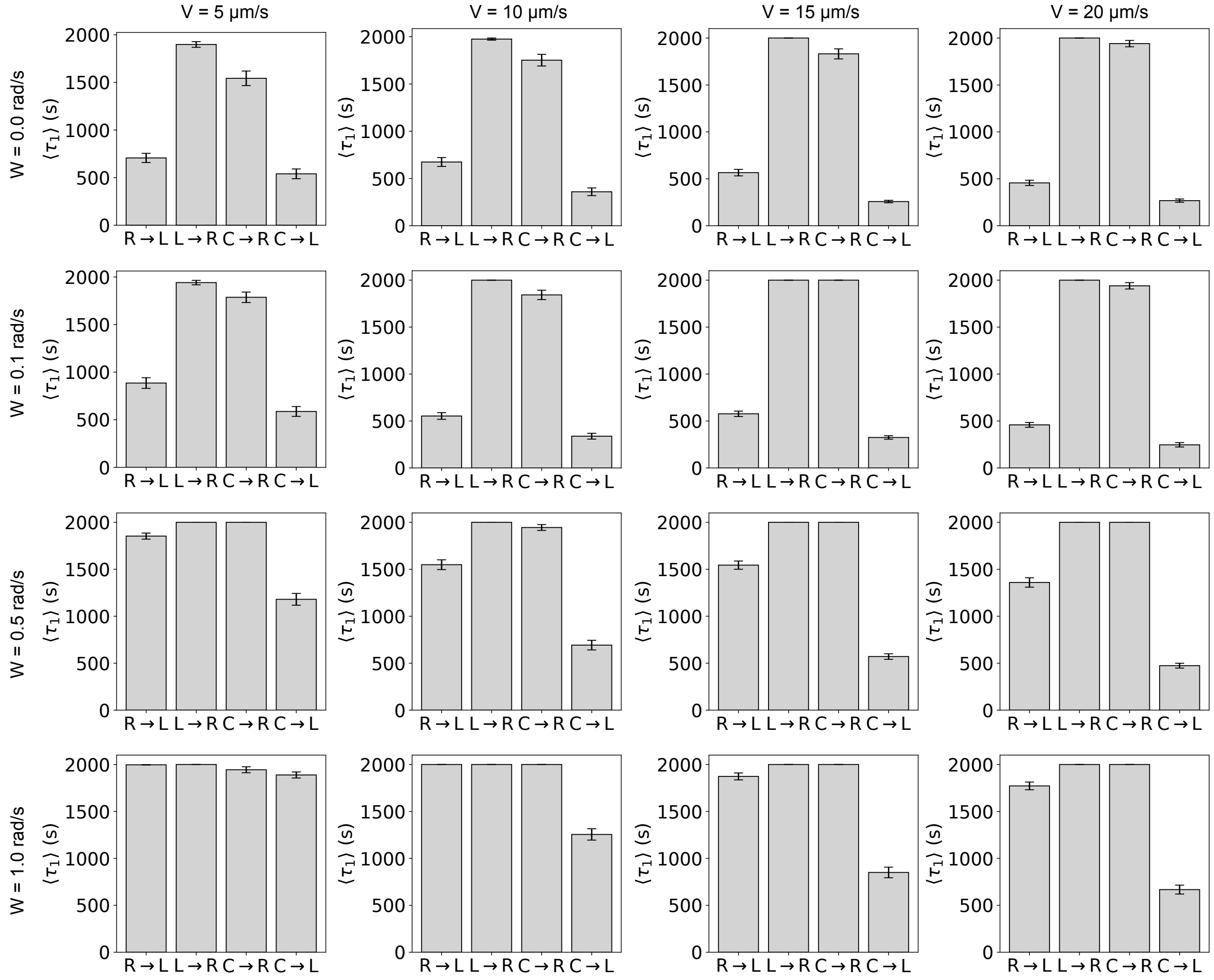


SI Figure 3: The average first passage time $\langle \tau_1 \rangle$ of active swimmers reaching the left or right pool with different initial positions, deterministic linear velocities ($V$, in $\mu$m/s), and angular velocities ($W$, in rad/s).
